# Effect of aging-related network changes on unimanual sensorimotor learning – a simultaneous EEG-fMRI study

**DOI:** 10.1101/2021.02.15.431203

**Authors:** Sabrina Chettouf, Paul Triebkorn, Andreas Daffertshofer, Petra Ritter

## Abstract

Sensorimotor coordination requires orchestrated network activity mediated by inter- and intra-hemispheric, excitatory and inhibitory neuronal interactions. Aging-related structural changes may alter these interactions. Disbalancing strength and timing of excitation and inhibition may limit motor performance. This is particularly true during motor coordination tasks that have to be learned through practice. To investigate this, we simultaneously acquired electroencephalography (EEG) and functional magnetic resonance imaging (fMRI) in two groups of healthy adults (young N=13: 20-25y and elderly N=14: 59-70y), while they were practicing a unimanual motor task. Both groups learned the task during brain scanning, which was confirmed by a 24h follow-up retention test. On average, quality of performance of older participants stayed significantly below that of the younger ones. Accompanying decreases in motor-event-related EEG-source beta band power (β, 15-30 Hz) were lateralized in both groups towards the contralateral side, albeit more so in younger participants. In the latter, the mean β-power during motor learning in bilateral pre-motor cortex (PM1) was significantly higher than in the older group. Combined EEG/fMRI analysis revealed positive correlations between fMRI signals and source-reconstructed β-amplitude time courses in contralateral and ipsilateral M1, and negative correlations in bilateral PM1 for both groups. The β-positive fMRI response in bilateral M1 might be explained by an increased cross-talk between hemispheres during periods of pronounced β-activity. During learning, the Rolandic β-power relative to rest was higher in bilateral PM1 in younger participants, suggesting less task-related beta band desynchronization in this (better performing) group. We also found positive correlations between Rolandic β-amplitude and fMRI-BOLD in bilateral M1 and negative correlations bilateral in PM1. This indicates that increased β-amplitudes are associated with increased M1 “activity” (positive BOLD response) and decreased PM1 “activity” (negative BOLD response). Our results point at decreased pre-motor inhibitory inputs to M1 as possible source for increased interhemispheric crosstalk and an aging-related decline in motor performance.

**Highlights:**

- Sensorimotor coordination performance decreases with increasing age.
- During motor learning the β-power in pre-motor areas is reduced with age.
- EEG/fMRI points at less effective inhibitory inputs from PM1 to ipsilateral M1 in older adults.

## 1. Introduction

Motor coordination requires a fine-tuned interplay of activities across the sensorimotor network (and beyond) that might be jeopardized by aging-related neurodegeneration (Holtrop et al., 2014; Sullivan and Pfefferbaum, 2006). Even in seemingly simple tasks like repetitive finger tapping, motor performance is known to decline with increasing age (Shimoyama et al., 1990) and that decline can be more pronounced when motor timing is more demanding or when movements are visually guided (Houx and Jolles, 1993; Kauranen and Vanharanta, 1996; Smith et al., 1999; Ward and Frackowiak, 2003). It might be that in older adults a proper motor control comes at the price of a widespread involvement of brain regions. Seidler et al. (2010) particularly highlighted prefrontal cortex and basal ganglia, which both are part of the motor network and are also known to be affected by aging-related alterations. Over the years, several studies addressed the effect of aging on the sensorimotor network activity, primarily affecting (bilateral) primary and premotor areas (Fujiyama et al., 2016; Maes et al., 2017; Seidler et al., 2010; Swinnen and Wenderoth, 2004). Functional magnetic resonance imaging (fMRI) with the blood oxygen level dependent (BOLD) contrast revealed that ageing is accompanied by less deactivation of ipsilateral primary motor areas and presumably there is also a reduced inter-hemispheric inhibition during unimanual movements (Coxon et al., 2010; Goble et al., 2010; Hinder et al., 2012; Levin et al., 2014; Van Impe et al., 2009), which may impair motor control (suppression) of homologous end-effectors (Hutchinson et al., 2002; Newton et al., 2005; Ward and Frackowiak, 2003). Functional MRI measures the blood oxygenation response to neuronal activity that is the result of a mixture of processes and hence an indirect measure of neuronal function that is difficult to interpret. So called fMRI “deactivations” and “activations”, that is task (or regressor) negative or positive correlations of fMRI do not simply reflect neuronal inhibition versus excitation – but in fact can be caused by both (Ritter and Villringer, 2002). Another challenge for the interpretation of fMRI is the low temporal resolution in the order of one or a few seconds typically. In contrast to fMRI, magneto- and electro-encephalography (M/EEG) can provide insight into the dynamics of learning a coordinated movement at high temporal resolution. Neural oscillation in different frequency bands are thought to mediate the functional coupling of the motor network. First and foremost the Rolandic α- and β-rhythms (around 8-14 and 15-30 Hz, respectively) are prominently present in sensorimotor regions (Pfurtscheller, 1981). While the mean β-activity decreases in the course of motor learning its motor-related modulation increases, suggesting its importance for motor timing (Boonstra et al., 2007; Houweling et al., 2010b). The abundant studies involving EEG have recently been summarized Rueda-Delgado et al. (2014), pointing at the pivotal role of the oscillation-based spatiotemporal reorganization of the motor network with demanding motor coordination.

Inter-limb coordination is a particular form of motor coordination, where left and right hemispheres have to orchestrate their activities. The prime interface for left/right interactions is the corpus callosum (CC), which is also known for its aging-related changes (Frederiksen and Waldemar, 2012; Langan et al., 2010; Sullivan et al., 2001; Sullivan and Pfefferbaum, 2006). In fact, the deterioration of CC may also affect uni-manual motor control, as there is accumulating evidence that proper unimanual performance entails the inhibition of ipsilateral motor areas (Chettouf et al., 2020; Daffertshofer et al., 2005; Ghacibeh et al., 2007; Gross et al., 2005; van Wijk et al., 2012; Vercauteren et al., 2008). Trans-callosal pathways are both inhibitory and excitatory (Daffertshofer et al., 2005; Swinnen, 2002). In particular the excitatory one may cause inter-hemispheric cross talk that may be visible in the form of (unwanted) mirror movements (Carson, 2005). Suppressing this cross talk hence requires intra-hemispheric inhibition in the hemisphere that is supposed to stay inactive. Yet, that inhibition needs to be timed, which might involve supplementary motor and/or premotor areas (SMA and PM1, respectively) as suggested by studies using transcranial magnetic stimulation (Stinear and Byblow, 2002). In a recent review we outlined a candidate scheme (Fig. 1) for the corresponding circuitry(Chettouf et al., 2020; Daffertshofer et al., 2005; Ghacibeh et al., 2007; Gross et al., 2005; van Wijk et al., 2012; Vercauteren et al., 2008), where during unimanual movements the contralateral M1 projects through the CC to excite its ipsilateral M1, while the same contralateral M1 also projects to the ipsilateral PM1. The latter has inhibitory cortico-cortical projections to M1 in the same hemisphere and, if properly timed, suppresses its activity stemming from contralateral M1. Disruption of PM1 activity by for example neurodegeneration in that area or in the CC, may impede this *effective inter-hemispheric inhibition*, hampering unimanual performance and/or cause activation of homologous muscles (Daffertshofer et al. (2005), or in the bimanual case, lead to instabilities of the planned motor action (Houweling et al., 2010a). With the current study we sought to investigate the proposed imbalance of the effective inter-hemispheric inhibition in the aging brain. This imbalance is expected to be particularly visible during learning a motor task with distinct timing requirements that we here used to contrast groups of young and older healthy adults.

**Figure 1:**
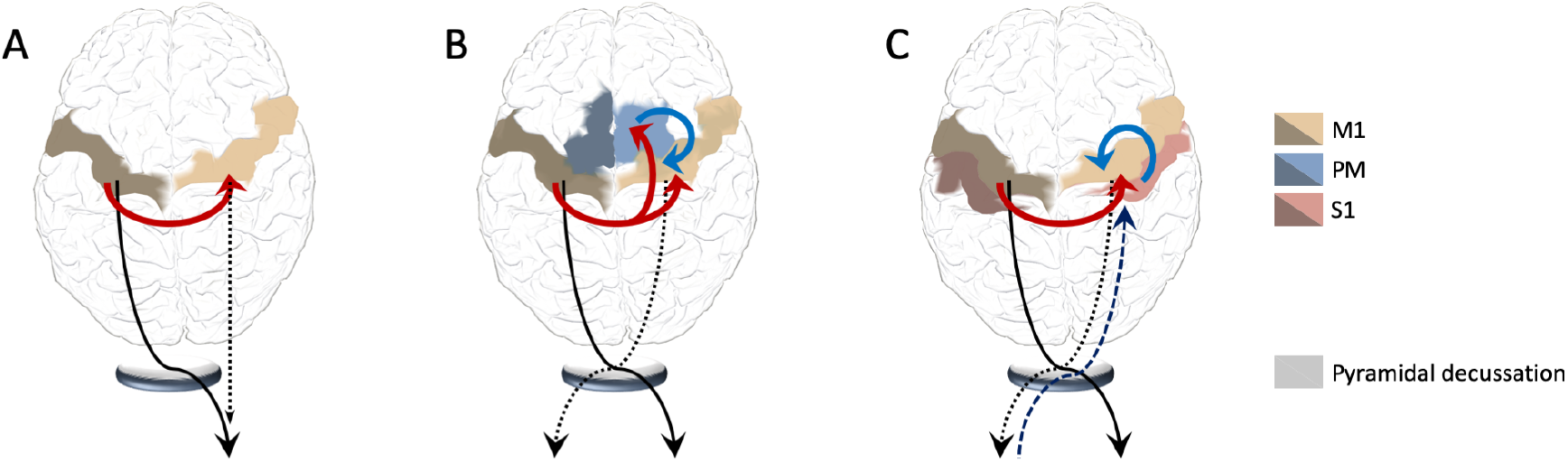
From Chettouf et al. (2020). Three alternative models that may account for bilateral cortical activation during unimanual performance. A: Left M1 is activated to generate motor output in a right end-effector, excitatory transcallossal connections activate right M1, which may contribute as ipsilateral controller. B: The model outlined in the *Introduction*. Left M1 activation causes a cross-talk through the corpus callosum in both, right M1 and right PM/SMA, with the latter inhibiting the first to prevent motor outflow to the left (homologous) end-effector. C: Again, left M1 activates right M1 but peripheral feed-back causes activation in right S1 that inhibits right M1 through cortico-cortical connections.

Every (brain imaging) modality comes with its own strengths and limitations. fMRI provides a spatial resolution in the order of cubic millimeters. On that aspect it can be considered superior to encephalographic, especially EEG, where neural source activity has to be inferred from surface recordings. However, fMRI relies on the assessment of large-scale metabolic and hemodynamic changes (quantified as BOLD signals) that are typically slow (~ seconds) and that do not directly capture neuronal (spiking) activity but is an indirect consequence of it. This neurovascular coupling acts as a low-pass filter of the underlying neuronal activity. That is, EEG provides a direct assessment of neuronal electric activity in up to the milli-second regime.

We hence combined both modalities, fMRI and EEG, in simultaneous recordings during a sensorimotor task (Ritter et al., 2009; Ritter and Villringer, 2006). First the recorded signals were analyzed separately following modality-specific standard approaches. The fMRI provided estimates of region-of-interests (ROIs) relevant for motor performance based on significant changes in BOLD signals. The EEG yielded motor-related activity patterns that, when source localized, revealed parts of the motor network that changed significantly in the course of learning and differed between age groups. Subsequently, the source-reconstructed EEG signals served as regressors to identify their correlates in fMRI-BOLD.

We expected both groups to learn the unimanual motor task but learning rate and overall performance to decrease with age. We hypothesized a less strongly inhibited ipsilateral M1 with increasing age in line with a reduced effective inter-hemispheric inhibition. That is, we expected 1) a less lateralized motor-related EEG beta-band modulation (indicating less interhemispheric feedforward inhibition) during the unimanual motor paradigm and 2) more task positive correlated activity in ipsilateral M1 (indicating more activation due to the lack of feedforward inhibition) in the older age group.

## 2. Material and methods

### 2.1 Participants

Twenty young adults (mean age = 22.0 years; range, 20–25 years; 13 females) and twenty old adults (mean age = 63.6; range = 59–70 years; 14 females) participated in the study. None of the participants had a history of neurological, psychiatric or chronic somatic diseases. None of them had musical background, which is known for altered motor timing and accompanying neural activity (Hughes and Franz, 2007). All participants were right-handed according to the Edinburg Handedness Inventory test (Oldfield, 1971) and the did not show any cognitive impairment assessed with the Montreal Cognitive Assessment (Nasreddine et al., 2005). They gave written informed consent prior to assessment. The experiments were performed in compliance with the relevant laws and institutional guidelines and were approved by the medical ethical committee of the Charité Medical Center in Berlin (EA1/060/14).

### 2.2 Experimental design

We adopted a motor learning protocol that was successfully implemented in previous MEG studies (Houweling et al., 2008; Houweling et al., 2010b). Participants had to learn a polyrhythmic motor task with a ‘simple’ perceptual goal. Providing the latter is known to accelerate motor learning (Mechsner et al., 2001), which renders it particularly useful for learning complex behavior in limited scanning time. We modified this protocol to unimanual motor learning in the MR scanner.

Participants were asked to squeeze in an air-filled rubber bulb with their right hand in a 4:3 frequency ratio to an external cue. The cue was provided visually by a disc that rotated at 1.8 Hz on a computer display (left). Squeezing the bulb let a second disc (right) rotate (Fig. 2). Their squeezing rhythm, however, was mapped in such a way that the two discs rotated at the same frequency, if the cue/performance ratio was 4:3. Put differently, participants had to perform rhythmic squeezing at a frequency of 1.35 Hz to achieve a 1:1 left/right disc rotation which can be consider the aforementioned, ‘simple’ perceptual goal.

**Figure 2:**
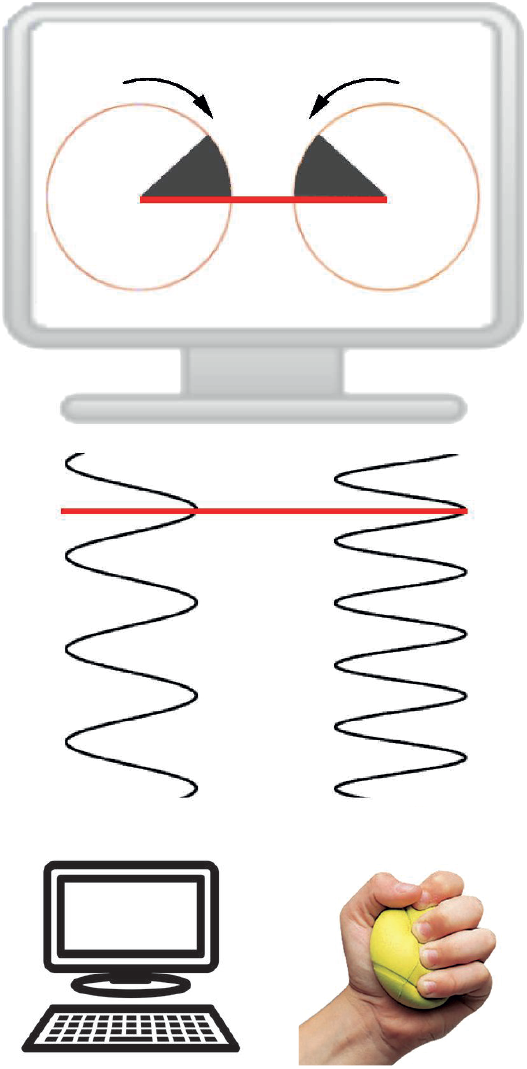
Experimental setup. The external cue and the force performed by the subject were displayed as two rotating discs on a computer screen that was projected via a mirror mounted on top of the head coil to the participant lying in the scanner. Squeezes of the rubber bulb with the right hand were analyzed in real time for the phase dynamics, which was multiplied by a factor 4/3. By this, proper performance of a 4:3 polyrhythm let the disks counter rotate with a 1:1 frequency ratio, which was typically realized in sync; Scheme has been modified from Houweling et al. (2008); Houweling et al. (2010b).

In the MR-scanner, participants performed a total of ten trials of two minutes each, separated by fifteen seconds breaks. They were instructed to keep their eyes open during the entire experiment and minimize eye- and head movements. The motor task was supplemented by two blocks of five minutes resting state recordings that served as baseline. There, participants were instructed to close their eyes, stay awake and to not move their head. On the consecutive day, participants performed the motor task outside the scanner, which we used as retention test for verifying motor learning rather than mere non-lasting training effects (Kantak and Winstein, 2012).

### 2.3 Data acquisition

Data have been acquired at the Berlin Center for Advanced Imaging at the Charité Universitätsmedizin Berlin.

#### 2.3.1 Motor behavior

We used a custom-made pressure transducer to convert the pressure inside an air-filled rubber bulb to an electrical signal that was sampled at a rate of 100 Hz (SCXI module, National Instruments, Austin, USA).

#### 2.3.2 EEG

EEG recordings were conducted with a 64-channel MR-compatible EEG system (Brain Products, Gilching, Germany; 0.1-250 Hz hardware band-pass filter, ±16.38 mV recording range at a 0.5 *μ*V resolution and 5 kHz sampling rate) and an EEG cap with ring-type sintered silver chloride electrodes with iron-free copper leads (EasyCap, Herrsching, Germany). 61 scalp-electrodes were arranged based on the international 10-20 system with FCz as reference and ground electrode at AFz. In addition, two electrocardiogram (ECG) electrodes and one vertical electro-oculogram (EOG) were recorded. An abrasive electrolyte gel (Abralyt 2000, Easycap, Herrsching, Germany) served to keep the impedances of all electrodes below 5 kΩ. The EEG-sampling was synchronized to the gradient-switching clock of the MR scanner, to ensure time-invariant sampling of the image acquisition artifact (SyncBox, Brain Products, Gilching, Germany) (Anami et al., 2003; Freyer et al., 2009).

#### 2.3.3 (f)MRI

We used a 3T Siemens Tim Trio MR scanner with a 12-channel Siemens head coil. Every scan session started with a localizer sequence (TR 20 ms, TE 5 ms, 3 slices (8 mm), voxel size 1.9×1.5×8.0 mm, FA 40°, FoV 280 mm, 192 matrix) followed by an anatomical T1-weighted scan (TR/TE 1900/2.52 ms, FA 9°, 192 sagittal slices (1.0 mm), voxel size 1×1×1 mm^3^, FoV 256 mm, 256 matrix), an anatomical T2-weighted scan (TR 2640 ms, TE1 11 ms, TE2 89 ms, 48 slices (3.0 mm), voxel size 0.9×0.9×3 mm, FoV 220 mm, 256 matrix). Afterwards, the EEG was prepared; fMRI (BOLD-sensitive, T2*-weighted, TR/TE 1940/30 ms, FA 78°, 32 transversal slices (3 mm), voxel size 3×3×3 mm, FoV 192 mm, 64 matrix) was recorded simultaneously to the EEG.

### 2.4 Data analysis

Data were analyzed using Matlab (version 2017b, The Mathworks, Natwick, MA). For the (f)MRI processing we also employed FreeSurfer (version 6.0.0, Laboratory for Computational Neuroimaging, Boston, US, see Fischl (2012)) the FMRIB (Functional MRI of the Brain) Software Library (version 5.0, Analysis Group, Oxford, UK, see Jenkinson et al. (2012)), connectome workbench (version 1.2.3, WU-Minn HCP Consortium, USA, see Marcus et al. (2011)) and the preprocessing scripts of the human connectome project (version 3.24.0, WU-Minn HCP Consortium, USA, Glasser et al. (2016); Glasser et al. (2013)).

#### 2.4.1 Preprocessing

EEG data were segmented with BrainVision Analyzer software (Brain Products) into three segments: the experimental learning task, pre- and post-resting state. Subsequently, we removed MR-scanner artifacts as detailed in the *Supplementary Materials*, down-sampled to 512 Hz and band-pass filtered between 0.1 and 100 Hz. Further EEG preprocessing consisted of removing and interpolating bad channels and removing the ballisto-cardiogram (pulsatile blood movement causing body and electrode movement inside the scanner) and excessive eye blinks as well as movement artifacts using independent component analysis with the fastICA package (Hyvarinen, 1999).

MRI data were preprocessed following the human connectome project preprocessing pipeline (Glasser et al., 2016; Glasser et al., 2013). Structural T1 and T2 weighted images were aligned, bias field corrected, skull stripped and nonlinearly registered to MNI space. We used the FreeSurfer recon-all to reconstruct cortical grey matter surfaces and a subcortical grey matter volume segmentation. Myelin maps across the surface were created by taking the ratio of T1w/T2w from voxel intensities inside the cortical grey matter (Glasser and Van Essen, 2011). FMRI data were motion corrected by aligning the first image to the series. Then, the motion parameters were used as regressors in the ICA denoising step to clean the fMRI data from remaining motion related artefacts. The fMRI data were EPI distortion corrected using a gradient echo field map, aligned to the T1w image as well as MNI space and bias field corrected. The fMRI data were converted into the CIFTI file format, where time series of voxels inside the cortical grey matter ribbon are mapped onto cortical surface vertices and the subcortical grey matter is resampled onto a standard volume mesh. Subsequently, fMRI data were cleaned using FSLs FIX tool (Griffanti et al., 2014; Salimi-Khorshidi et al., 2014). In brief, using a pre-trained classifier automatically we labeled independent components of the data as neural activity or artefacts. Then, motion parameters and artefact components served as regressors to remove corresponding variance from the fMRI data. Finally, the FIX classifier was trained on hand-labelled data from 23 subjects of this study. Subjects individual cortical surfaces were registered to the parcellation template (Glasser et al., 2016) of the human connectome project using a multimodal surface registration approach (Robinson et al., 2018; Robinson et al., 2014). The features used for the surface registration were cortical folding, myelin maps, resting state network locations and visuotopic maps. The pre- and post-task fMRI data were used to identify subject individual resting state network locations and visuotopic maps. Aligned fMRI data entered statistical analysis.

#### 2.4.2 Motor behavior

We quantified performance via the strength of frequency locking between the visual cue and the force produced. As corresponding measure we utilized normalized spectral overlap (Daffertshofer et al., 2000), i.e. the similarity 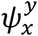 between the power spectrum of the cueing signal, *P*_*x*_, and that of the produced force, *P*_*y*_, after rescaling the frequency axis by the factor *ρ* = *p*: *q*. This measure reads

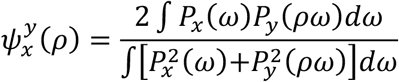

and is bounded to the interval [0,1] with 1 indicating maximum similarity. As common, we Fisher-transformed this value prior to statistical evaluation to stabilized normality.

Statistical analysis of the behavioral data was performed with SPSS (IBM Corp. Released 2015. IBM SPSS Statistics for Macintosh, Version 23.0 Armonk, NY: IBM Corp.). Frequency locking values per trial for both groups were subjected to an ANOVA with repeated measures. Within-subject factors were the ten task trials and the retention test and age were included as between-subject factor. The significance threshold was set to α = 0.05. Sphericity was tested using Mauchly’s test and, when violated, we applied a Greenhouse-Geisser’s correction. Post-hoc paired-samples t-tests were performed whenever a main effect of trial or age was detected to evaluate effects of motor learning.

#### 2.4.3 EEG

Source localization of the selected EEG rhythm was realized using dynamical imaging of coherent sources beamformers (Gross et al., 2005) with tissue segments (skin, skull, cortical spinal fluid, gray and white matter) determined through finite-element-modeling. For both we employed the open-source Fieldtrip toolbox and used their template MRI (Oostenveld et al., 2011).

##### Beamformers for motor event-related β-power

We adopted a motor-related approach with maximum changes of hand force as central events. Epochs of 400ms before and after these events served as contrast for subsequent statistics. To identify significance of beamformer power we followed the Monte Carlo approach with cluster-based test statistics for both groups (with a significance threshold of α_cluster_ = 0.01, Nichols and Holmes (2002)). Cluster-level statistics were determined as the sum of *t*-values per cluster. Probabilities were determined by collecting trials of pre- and post-event intervals and determined test statistics on random partitions. These steps were repeated 8,192 times to construct the permutation density of the test statistics, which allowed for using a dependent samples *t*-test between pre- and post-event epochs (significance threshold α = 0.05). The findings for the Rolandic α-frequency band can be found as *Supplementary Material*. Beamformer outcomes were parcellated using the SPM anatomical atlas (Eickhoff et al., 2005). Based on these outcomes, virtual sensors were defined for the ROIs onto which the EEG data were projected after band-pass filtering in the frequency band under study. The motor-event-related β-amplitude modulation per trial and for the two resting state blocks was evaluated as described in Houweling et al. (2010a). In brief, we normalized the β-band time-series of the virtual sensor to baseline (the first resting state recording), computed the mean Hilbert amplitude of this normalized data and performed a time-locked averaging over the aforementioned ±400 ms epochs.

##### Beamformers for task-vs-rest β-power

We also computed grand-average source-level β-power across learning versus pre-learning rest for both groups. We tested whether task modulations differed between groups using a voxel-wise permutation tests with an independent samples t-test.

#### 2.4.4 EEG regressors for analysis with fMRI

We analyzed the fMRI data using general linear modelling (GLM) as outlined below. When combining fMRI with EEG, the source localized EEG served as an additional regressor. For this, we estimated the instantaneous Hilbert amplitude in the frequency-band of interest and averaged it for every scan epoch (700 in total). To accommodate for delays and dispersions in the BOLD-responses, these regressors were convolved with the double gamma hemodynamic response function (HRF) provided by the FMRIB Software Library; see Fig. 2.

#### 2.4.5 fMRI/EEG

Before statistical analysis, the fMRI data were spatially smoothed both on the surface and in the subcortical volume using a 4 mm FWHM Gaussian Kernel to suppress spatial noise and increase the signal-to-noise ratio. To remove low frequency noise, the data were also filtered with a Gaussian-weighted linear high-pass filter with a cutoff of 135s. On single-subject level, we fitted the GLM: *Y* = *β*_*k*_*X*_*k*_ + *ε*, using the grayordinates-wise BOLD data *Y* in CIFTI format, i.e. the time series from cortical vertices and sub-cortical voxels (Friston et al., 1995). Here, *β*_*k*_*X*_*k*_ denotes the design matrix and *ε* the residual error. To build the design matrix we used the task paradigm and also combined it with the EEG source-localized β-power (Fig 3, bottom panel). The latter stemmed either from contralateral M1 (left area 4a) or from frontal cortex (left area 6 ~ premotor area), both according to the SPM atlas (Eickhoff et al., 2005). We contrasted each regressor against baseline and both against each other to estimate statistical effects of interest. When both regressors - task and EEG – are placed in a single GLM, the parameter estimates will reflect the activation in BOLD of one regressor adjusted for the effect of the other. In consequence the variance explained by both regressors will be removed, isolating the effect of, in our case, the EEG regressor. When using orthogonalization, the shared information of the two regressors is attributed to the regressor that is not orthogonalized (Mumford et al., 2015). Hence, we did not apply any orthogonalization. Based on these single-subject contrasts, a mixed-effects group-level analysis was performed using an unpaired two group t-test, to contrast within- and between-group activation. The latter contrasts were masked to reveal only those areas that deemed significant using the single-group activation contrast. Finally, we thresholded the statistical maps using the false discovery rate of *q* = 0.01 (Genovese et al., 2002).

**Figure 3:**
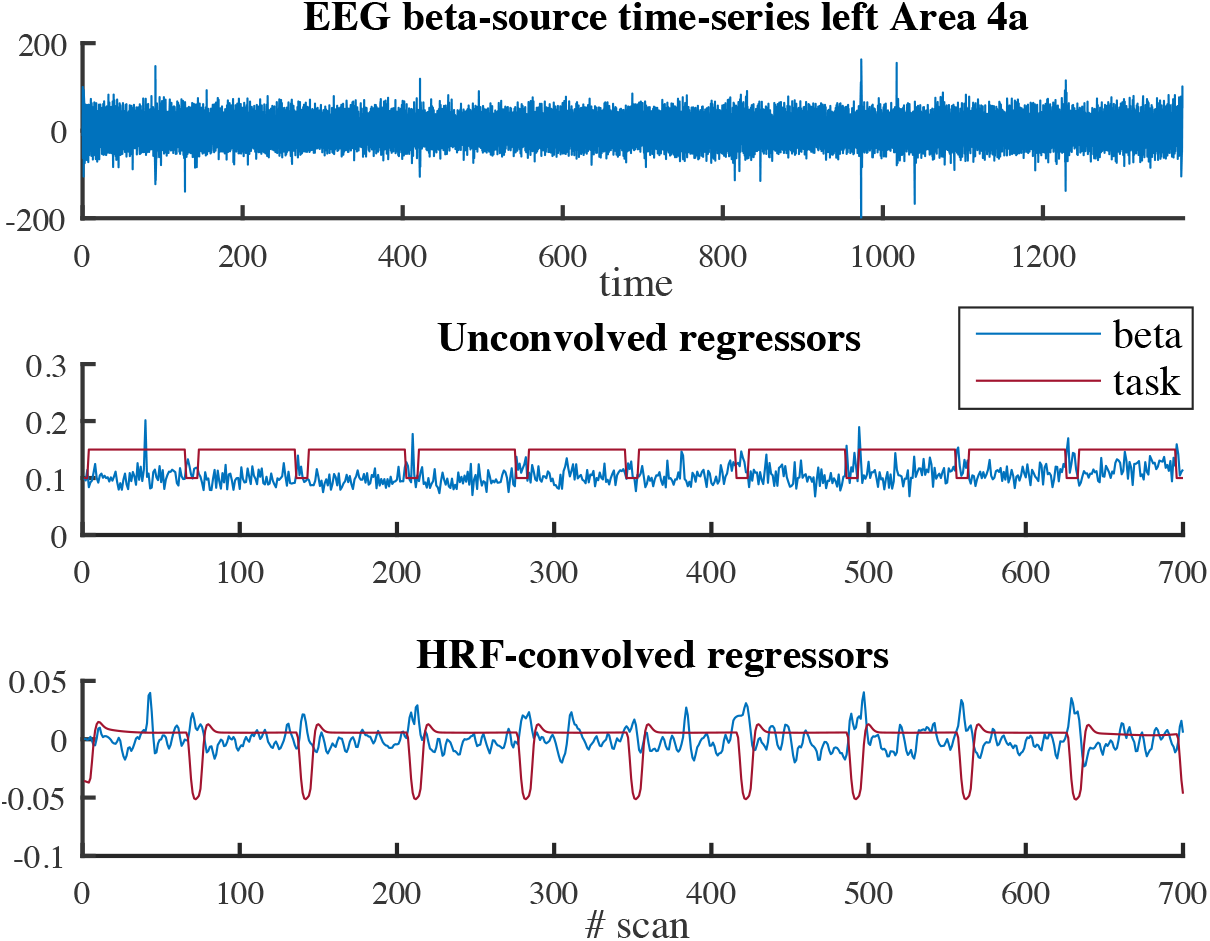
Computation regressors. Illustration of the computation of the beta amplitude and the resulting regressors. From the source localized EEG in the beta band (top panel), the Hilbert amplitude of every scan was determined (middle panel, blue) and convolved with the hemodynamic response function (HRF) to determine the HRF-convolved regressor (bottom panel, blue). The red curves show the regressor of the task design (in the bottom panel after HRF-convolution).

**Figure 4:**
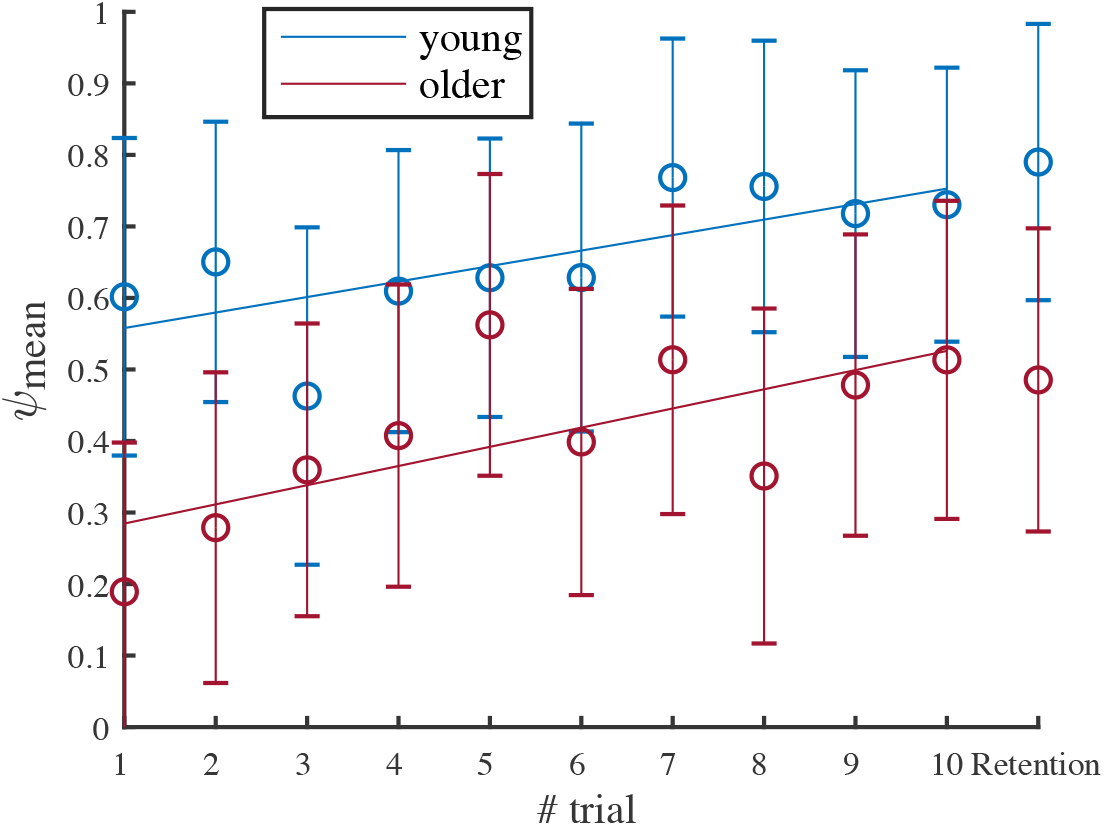
Behavioral results. Mean performance and standard error per trial for 13 young (blue) and 14 older (red) participants during the fMRI/EEG recordings and the mean performance and standard error of the retention test.

## 3. Results

EEG data of eleven participants (six young, five older) had to be excluded due to technical errors during acquisition. Two subjects showed extensive head motion and had to be excluded from further fMRI analysis. Data of 27 participants entered behavioral, EEG and fMRI analysis.

### 3.1 Motor behavior

Since the assumption of sphericity was violated (*χ*^2^(44) = 83.43, *p* < .001), we corrected the degrees-of-freedom (*ε*= .564). We found significant main effects for trial (*F*(5.08,126.99) = 3.621, *p* < .01) and age (*F*(1,25) = 11.723, *p* < .01) but could not establish a significant age × trial interaction. The performance level during the first trial was significantly lower in the older than in the young participants (*p* < 0.05). A difference that did not persist during the last trial, but again present during the retention test.

Post-hoc t-tests revealed an improved performance during trial 10 when compared to trial 1 in the older group (*p* = .004), but not for the younger group (*p* = .072). Both groups showed significant differences between the first trial and the retention test (young: *p* < .05, older: *p* < .01), while we could not identify any significant differences between the last trial and the retention test (*p* > .05).

### 3.2 EEG

As summarized in Fig. 5, the β-band power that contrasted 400ms pre- and post-motor event displayed significant motor-task positive modulation (i.e. an increase during the motor event) that was lateralized in both groups. Differences between groups over the complete task were not significant. We used left area 4a, i.e. M1 contralateral to the force producing hand, as ROI for subsequent analyses because, on group level, it displayed the maximum absolute *t*-value: *t* = −6.94 (*p* < .05) in younger and *t* = −3.73 (*p* < .05) in the older group.

**Figure 5:**
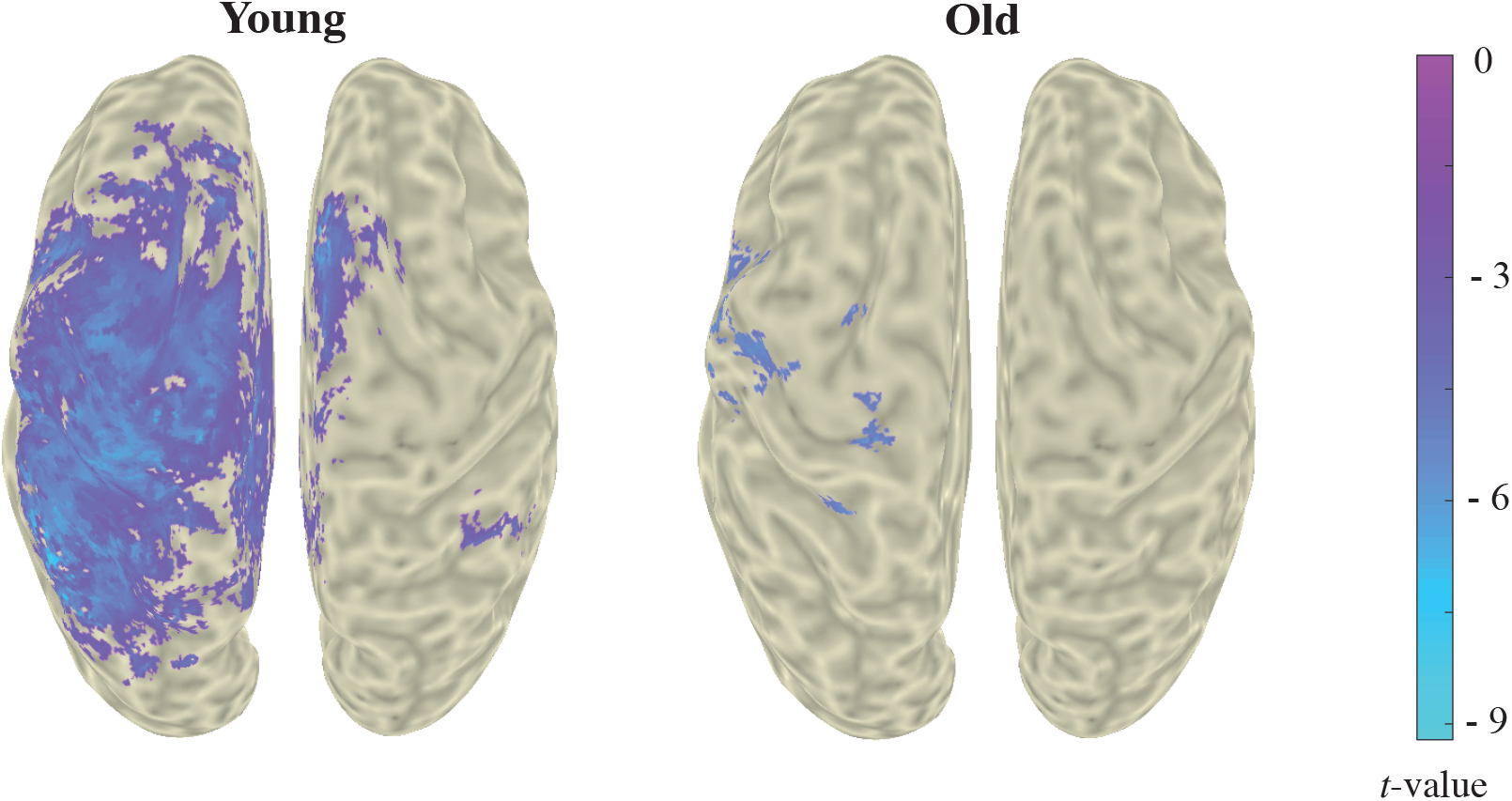
Event-related beamformer results for the β-band power. Adaptive spatial filtering identified cortical sources where β-band power correlated positively with the motor task execution. Grand averages shown correspond to young (N=13; left) and old (N=14, right) participants. Permutation tests with a dependent samples t-test as test statistic revealed voxels comprising significant differences between 400ms pre- and post-motor event for both young and older groups. Colors represent *t*-values masked with a threshold of *p* < .05 for the young and older groups.

For both groups, we found motor-event-related β-amplitude increases in contralateral M1, which was more pronounced in the young group (Fig. 6a). However, the standard deviation – in light blue for young and red for old – shows a great intra-group variability and therefore no group differences can be found for this event-related modulation. Yet, when we do not normalize the data to the average β-amplitude during the first baseline resting state recording, and thereby investigating the beta amplitude modulation between pre-learning resting state and learning, we see - besides a drop from rest to task - an overall lower mean beta power for the old group compared to the young during the motor task (Fig. 6b). This is in line with the results shown in Fig. 7, where the groups are compared by using a grand average of all task blocks. The figure displays were the young reveal a higher mean beta power compared to the older group, with a peak *t-*value of 4.14 (*p* < .05) in left area 6 (premotor area).

**Figure 6a:**
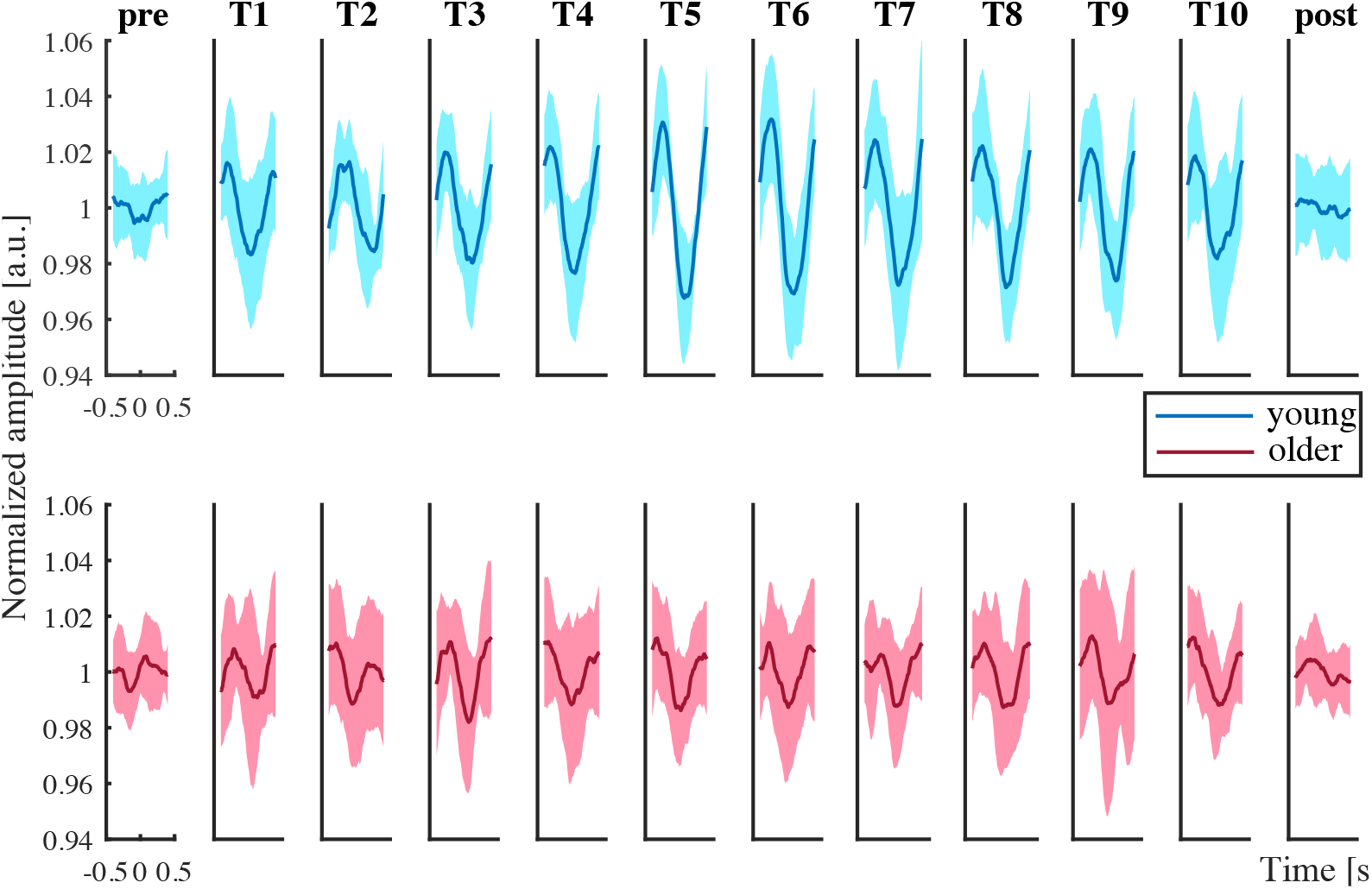
Learning block-wise event-related β-amplitudes in contralateral (left) M1. Grand average of event-related amplitudes in left M1 (area 4a) for pre- and post-rest and ten task blocks for the young (upper row in blue, N=13) and older (lower row in red, N=14) group; amplitudes (in arbitrary units, including standard deviations) are normalized to the mean β-amplitude value of the first resting state trial (baseline).

**Figure 6b:**
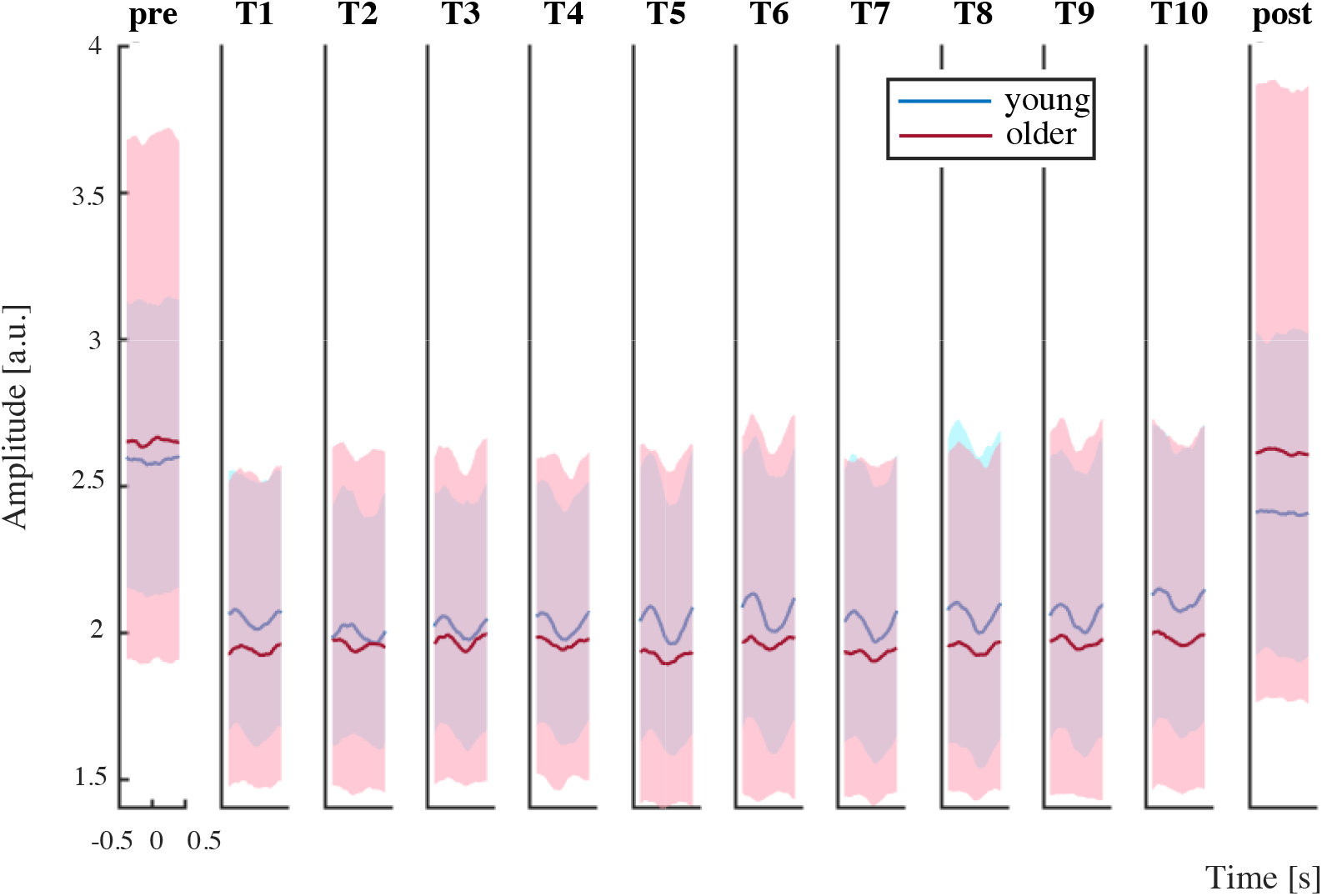
Non-normalized event-related β-amplitudes in contralateral (left) M1. Grand average of event-related amplitudes in left M1 (area 4a) for pre- and post-rest and ten task blocks for young (blue, N=13) and older (red, N=14) group; amplitude and standard deviations are displayed in arbitrary units.

**Figure 7:**
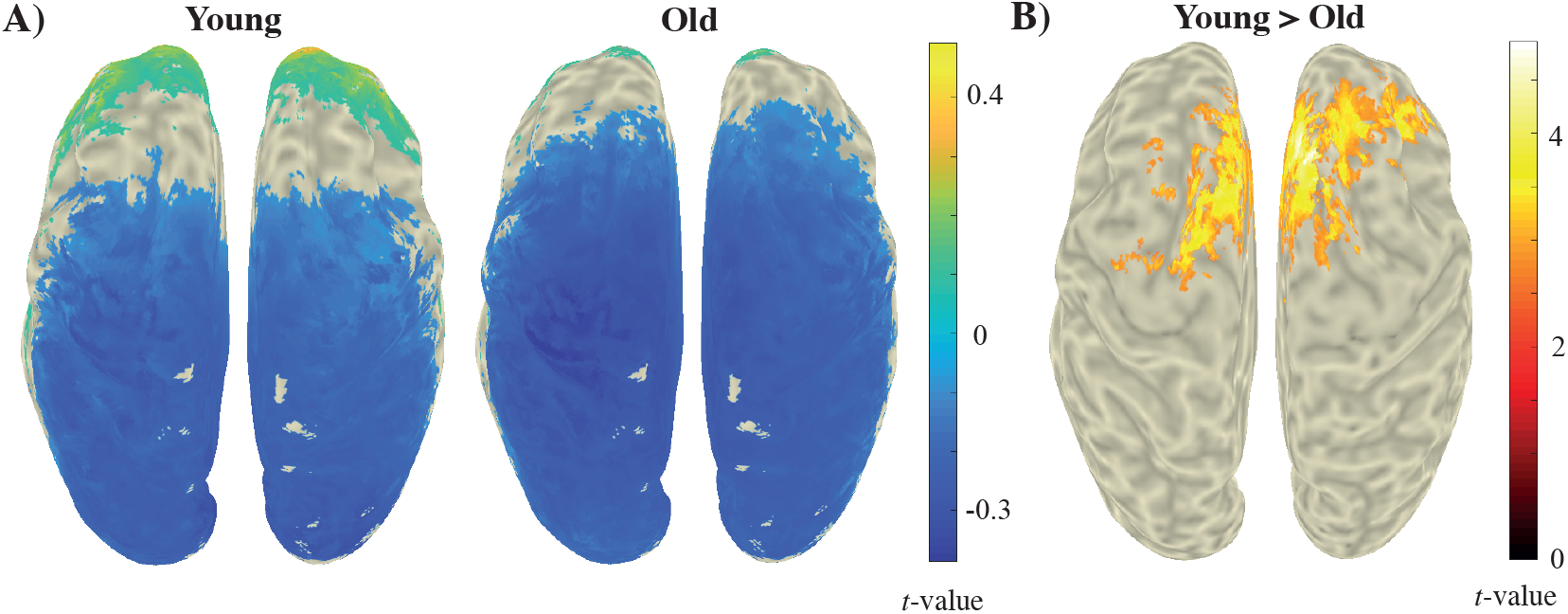
Grand averaged differences of β-power learning-task. A) Task-versus-rest descriptive statistics of β-power decreases for both age groups. B) Permutation tests with an independent samples t-test as test statistic revealed voxels comprising significant β-power differences during learning between young and older groups. Colors represent *t*-values, masked with a threshold of *p* < .05.

### 3.3 fMRI

We observed significant task-induced BOLD signal modulations in a number of visual and motor regions both in young and older participants (FDR corrected *p* < .01). As expected, the right-hand motor task was primarily accompanied with activity in left M1. We also found task-related activity in left and right PM1 in both groups and in SMA in the young one (Fig. 8). And, subcortical areas were active that are known for being involved in regulating movement and motor learning, including putamen, thalamus and cerebellum.

**Figure 8:**
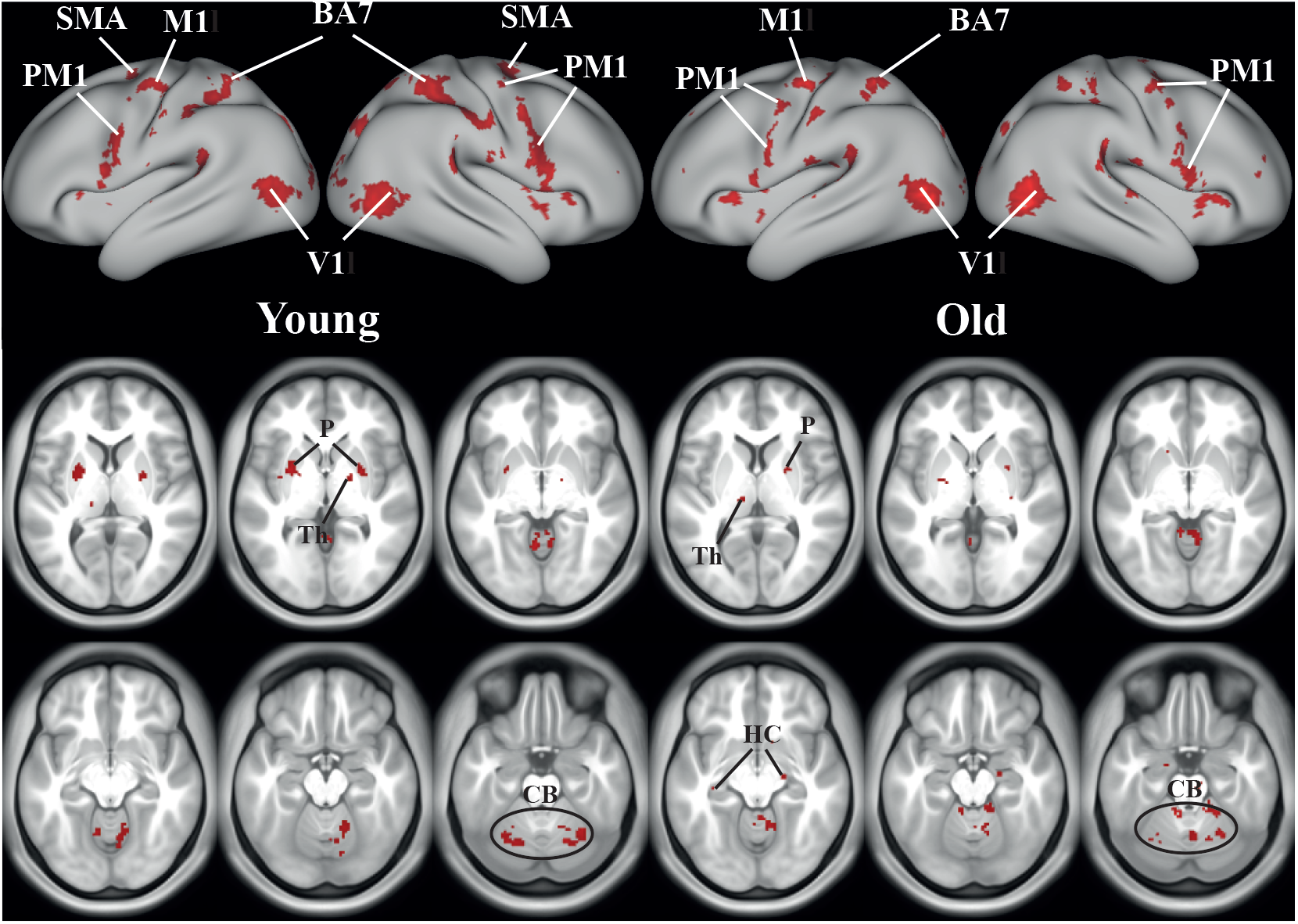
Task-positive BOLD responses. Using the task design as regressor, we identified significant clusters of task positive BOLD responses displayed in red for young (left, N=13) and older (right, N=14) participants (unpaired two group t-test, FDR-corrected *p* < .01). We used the Glasser parcellated brain template to display the BOLD patterns. PM1 = premotor cortex; M1 = motor cortex; SMA = supplementary motor cortex; BA7 = Brodmann area 7; V1 = visual cortex; P = putamen; CB = cerebellum; Th = thalamus.

To identify brain areas with task-related activity, we used the task design as regressor, yielding patterns in both groups that largely resemble the default mode network (DMN, see Fig. 9). According to Andrews-Hanna et al. (2014), the DMN consists of the posterior and anterior cortical midline structures, with hubs located in the posterior cingulate cortex and precuneus, the medial prefrontal and the parietal cortex (Brodmann areas 7 and 39/40). We did not find significant young versus older group differences, neither in task positive nor negative BOLD responses during motor learning.

**Figure 9:**
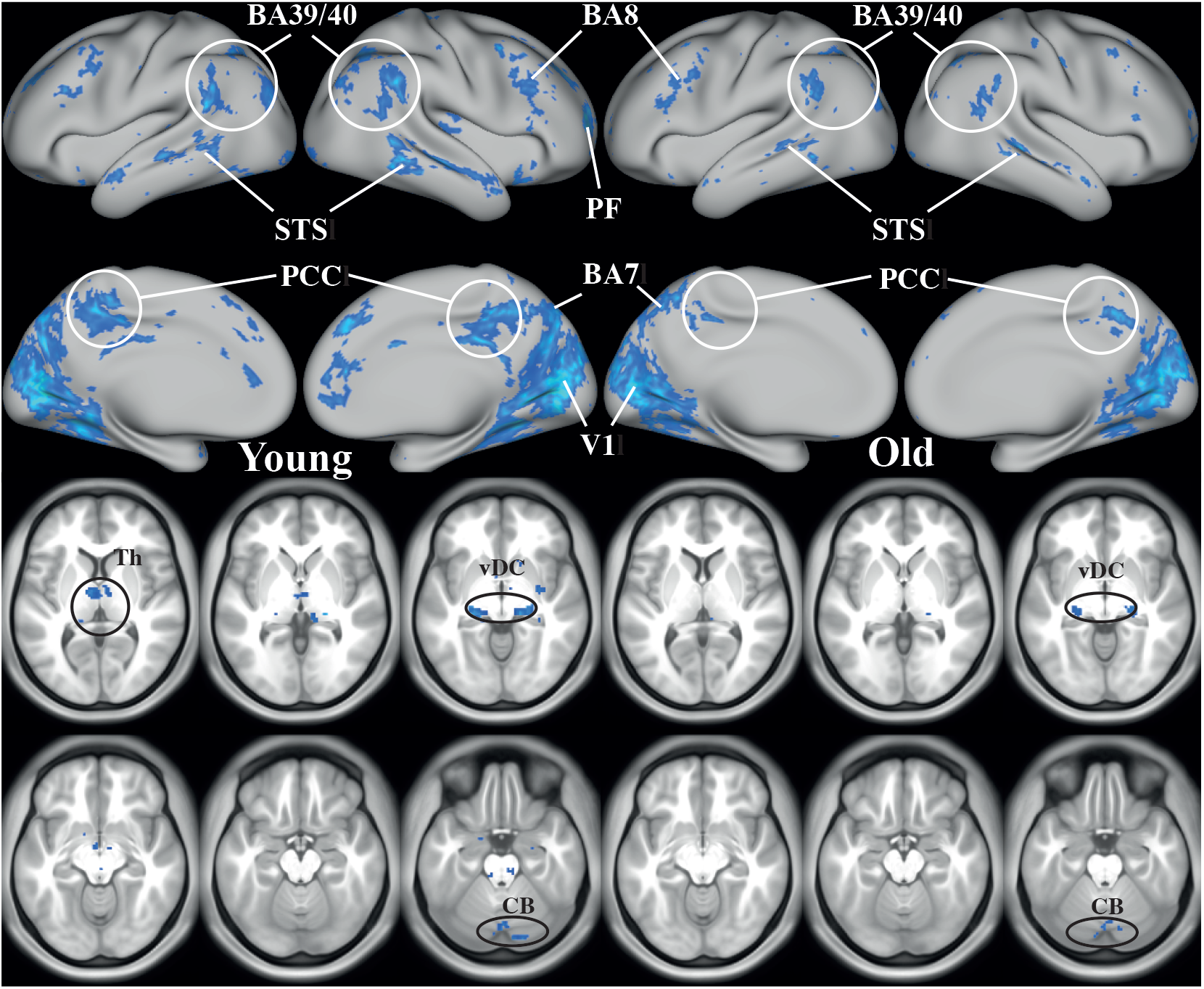
Task-negative BOLD responses. The task design as regressor revealed regions that significantly deactivated in blue for young (left, N=13) and older (right, N=14) participants (unpaired two group t-test, FDR-corrected *p* < .01). BA7/8/39/40 = Brodmann area 7/8/39/40; PF = prefrontal cortex; STS = superior temporal sulcus; PCC= posterior cingulate cortex; V1 = visual cortex; CB = cerebellum; Th = thalamus; vDC = ventral diencephalon.

### 3.4 fMRI/EEG

Since the EEG beamformer results revealed left M1 as main ROI we employed the corresponding β-amplitudes to define supplementary regressors. By this we expected to unravel regions related to modulation in β-activity. For both groups, the regions that significantly correlated positively with the β-amplitude regressor were primary motor- and visual cortex, Brodmann’s area 9, 39 and 32, putamen and hippocampus. The activated Brodmann areas are considered to contribute to short-term memory and attention (Fig. 10). For the older adults also the primary somatosensory cortex, insula, thalamus and amygdala came significantly to the fore. The insula is known for its involvement in hand-and-eye motor movement and motor learning. A remarkable difference in comparison to the task-correlated fMRI patterns, is a positive correlation in the right (ipsilateral) M1 present both groups. Moreover, bilateral premotor areas, the anterior intraparietal area and some visual areas showed a significant negative correlation when including the β-amplitude regressor (Fig. 11). And, the older group showed a negative correlation between fMRI-BOLD and beta amplitude in M1 in a small part of the left M1. Again, we could not find any significant group difference.

**Figure 10:**
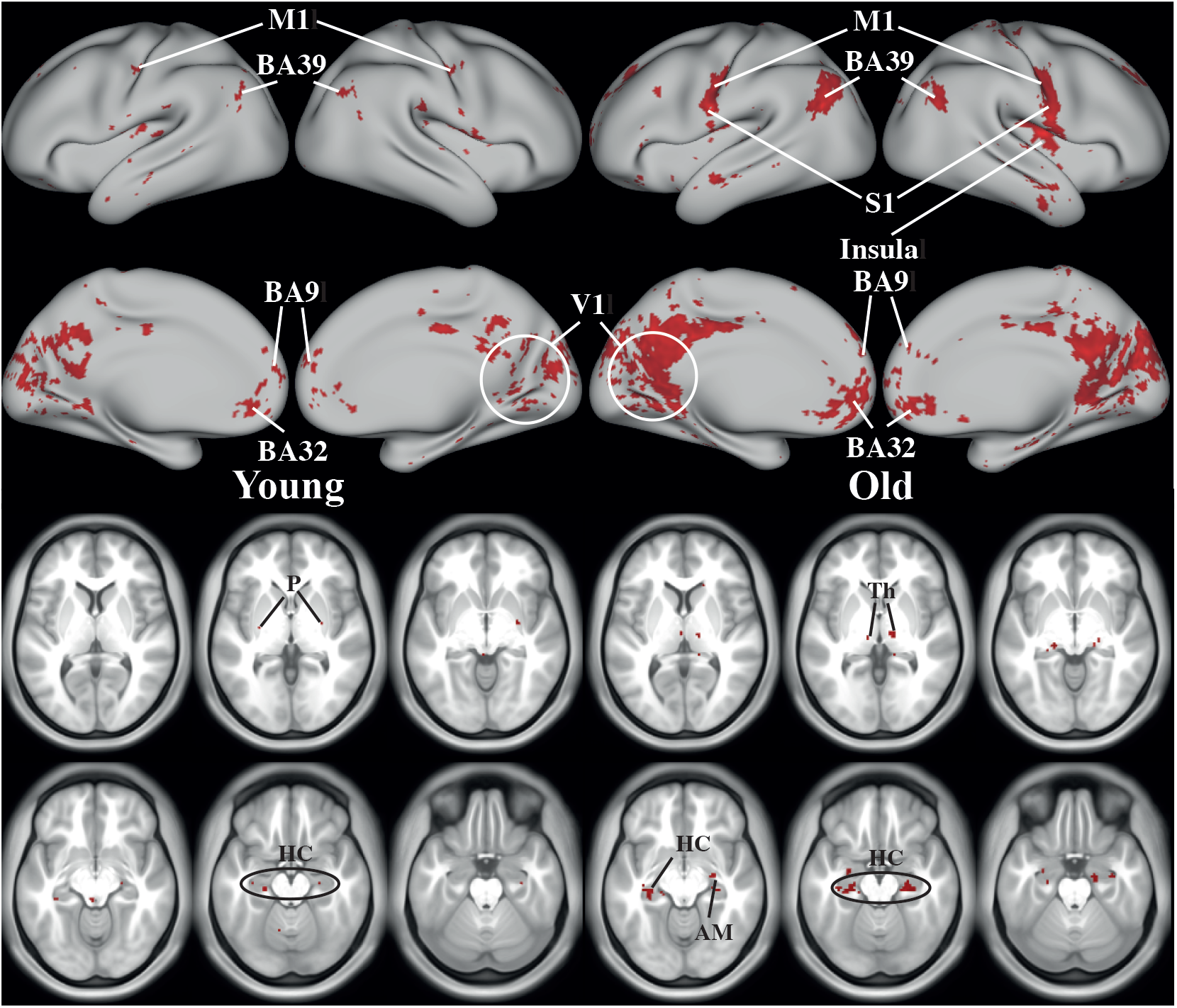
BOLD responses when including β-amplitudes in left M1 as additional regressor. Using both the task design and motor-related β-amplitudes in left M1 as regressors, we evaluated regions that significantly correlated in red for young (left, N=13) and older (right, N=14) participants (unpaired two group t-test, FDR-corrected *p* < .05). M1 = motor cortex; S1 = somatosensory cortex; BA9/32/39 = Brodmann area 9/32/39; V1 = visual cortex; P = putamen; CB = cerebellum; Th = thalamus; AM = amygdala; HC = hippocampus.

**Figure 11:**
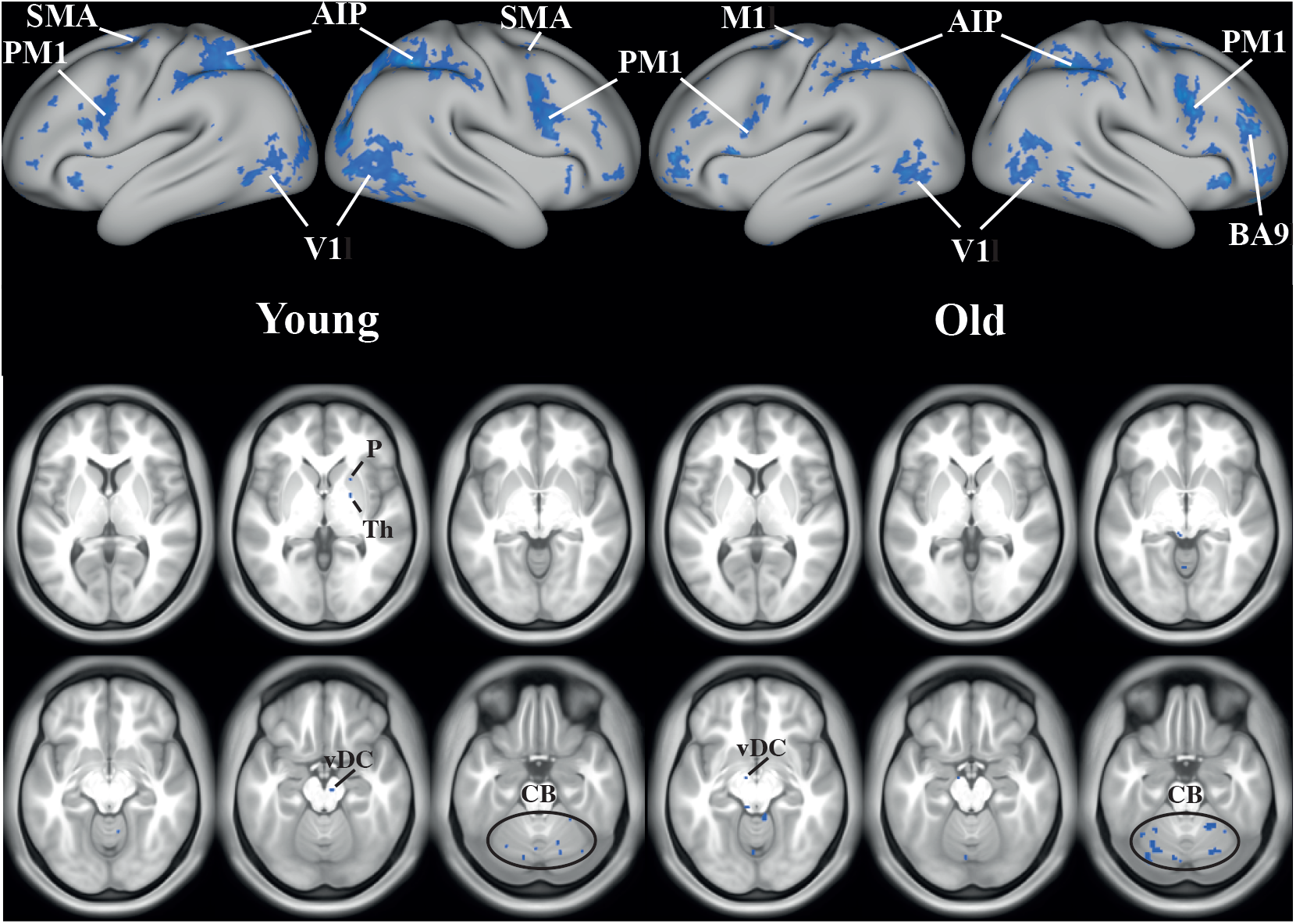
Negative fMRI-BOLD signal correlates of β-amplitude in M1. Regions that significantly correlated negatively are indicated in blue for young (left, N=13) and older (right, N=14) participants (unpaired two group t-test, FDR-corrected *p* < .05). PM1 = premotor cortex; M1 = motor cortex; AIP = anterior intraparietal area; SMA = supplementary motor cortex; BA9 = Brodmann area 9; V1 = visual cortex; CB = cerebellum; Th = thalamus.; vDC = ventral diencephalon.

Recall that we did find a significant group difference (young > old) in the mean β-power; cf. Fig. 7. There, Brodmann’s area 6 turned out to be the most significant ROI. Therefore, we used the corresponding source-reconstructed β-amplitude as regressor for the BOLD analysis. This revealed regions similar to the afore-described analysis using the β-amplitude in left M1, albeit the regions appeared more focal (Fig. 12). Still, group differences in fMRI-BOLD could not be established despite the differences in EEG.

**Figure 12:**
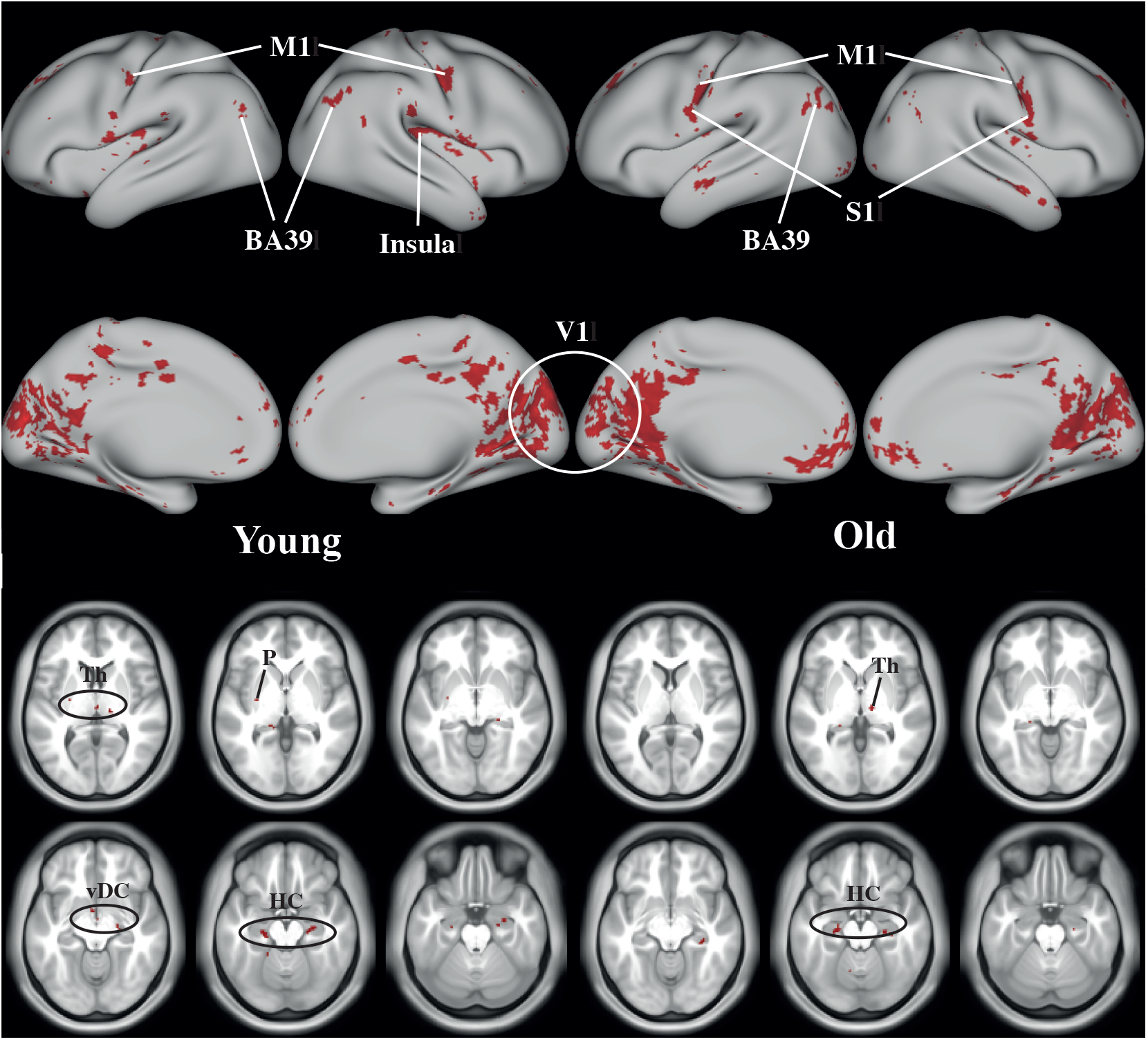
BOLD responses based on β-amplitude in left area 6 as supplementary regressor. With task design and the motor-related β-amplitudes in left area 6 (left PM) as regressor, we found several regions that significantly correlated in red for young (left, N=13) and older (right, N=14) participants (unpaired two group t-test, FDR-corrected *p* < .05): M1 = motor cortex; S1 = somatosensory cortex; BA39 = Brodmann area 39; V1 = visual cortex; P = putamen; Th = thalamus; vDC = ventral diencephalon; HC = hippocampus.

**Figure 13:**
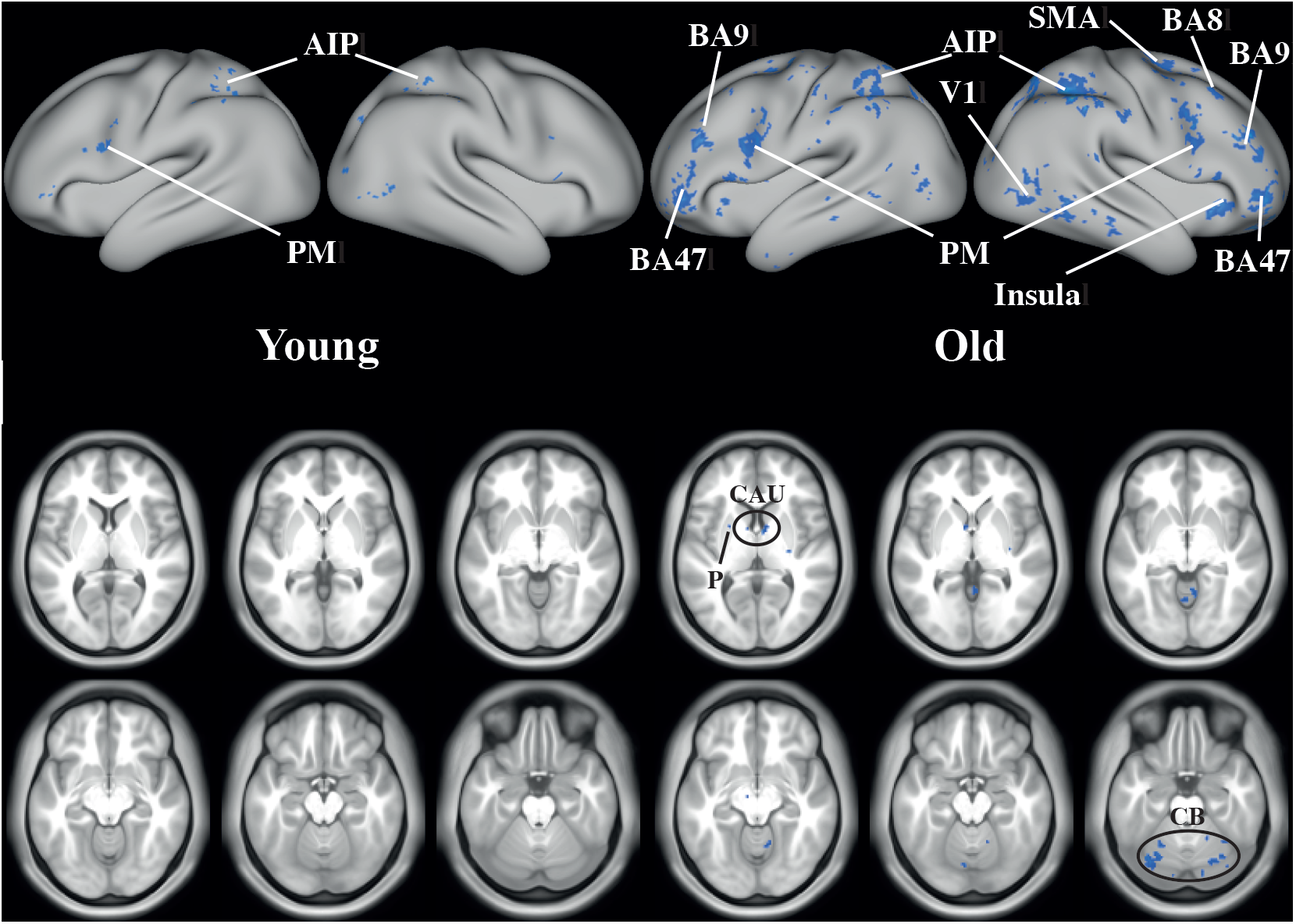
BOLD negative responses using the β-amplitude in contralateral area 6 as supplementary regressor. Several regions were significantly deactivated in blue for young (left, N=13) and older (right, N=14) participants (unpaired two group t-test, FDR-corrected *p* < .05). PM = premotor cortex; AIP = anterior intraparietal area; SMA = supplementary motor cortex; BA8/9/47 = Brodmann area 8/9/47; V1 = visual cortex; CB = cerebellum; P = putamen; CAU = caudate.

## 4. Discussion

We simultaneously acquired EEG and fMRI-BOLD signals in younger and older participants during the performance of a unimanual motor learning task. The experiment was designed to unravel aging-related differences in motor coordination learning. Both age improved performance due to learning. The training was accompanied by task-related changes in the motor event-related EEG β-power (Fig. 5) and in the mean β-power over all task blocks (Fig. 7). The latter differed significantly between groups in premotor areas. The fMRI-BOLD also revealed significant changes in the course of the motor task, but in contrast to EEG it did not reveal any significant differences between younger and older participants.

Based on the behavioral outcomes, both groups were able to learn the motor task to a similar degree. Yet, there was an offset implying poorer performance in the older group that persisted throughout the learning. When correcting for this by stratifying the starting performance level, the rate of learning largely agreed between groups in line with earlier findings (Bhakuni and Mutha, 2015; Hoff et al., 2015). The 24-hours follow-up retention test confirmed motor learning in both groups rather than mere training effects. That is, as expected also during the retention test we observed decrease in motor performance with increasing age (Houx and Jolles, 1993; Kauranen and Vanharanta, 1996; Shimoyama et al., 1990; Smith et al., 1999; Ward and Frackowiak, 2003).

The β-band beamformers were signified by activity in left M1, i.e. contralateral to the force producing hand. This was the case in both groups albeit stronger in the younger one. We expected to find less lateralized activity in the older adults than in the young one (Cabeza, 2001) since – by hypothesis – aging may limit the deactivation of the ipsilateral M1 during unimanual movement because of a reduced increase intrahemispheric inhibition due to compromised phase locking between premotor and primary motor areas (Coxon et al., 2010; Daffertshofer et al., 2005; Goble et al., 2010; Hinder et al., 2012; Van Impe et al., 2009; van Wijk et al., 2012). Yet, we could not confirm this expectation as ipsilateral activity did not reach statistical significance, possibly due to a substantial within-group variability which is known to be present in transient motor behavior, here motor learning (Bernstein, 1967; Riley and Turvey, 2002). This variability may have also caused the lack of significant group differences in motor-related β-modulation in M1 (Fig. 6a). However, we found significant differences between the groups in mean β-power located in (bilateral) premotor areas. Unfortunately, this camouflaged possible learning-related differences over time as all task block were combined in this statistical comparison. Yet it is striking that the task positive β-power in the younger participants was substantially elevated when compared to the older ones in prefrontal regions. In fact, this overall decrease in β-power when contrasting an uni-manual motor task with rest is well described in EEG/MEG research (Calmels et al., 2008; Calmels et al., 2006; Farber and Anisimova, 2000; Ford et al., 1986; Gerloff et al., 1998; Lange et al., 2006; Man’kovskaya, 2006; Manganotti et al., 1998; Mima et al., 2000; Serrien, 2008; Shibata et al., 1998; Svoboda et al., 2002; van Wijk et al., 2012; Vecchio et al., 2014; Wang et al., 2017). Moreover, the coherence between bilateral premotor and sensorimotor areas appears to increase. The results of the current study show that the drop in β-power is significantly greater in older compared to younger adults, with premotor areas as main region with age-related differences. This is in line with Espenhahn et al. (2019), who showed that the magnitude of movement-related beta desynchronization is affected by age: older subjects showed a greater β-power decrease in both sensorimotor cortices during the movement than their younger counterparts.

The fMRI-BOLD patterns contained significant task-related activation also in left M1, in bilateral PM1, SMA and visual areas, while the deactivation patterns largely agreed with the DMN. None of these differed significantly between age groups. The lack of aging-effects in fMRI-BOLD persisted despite supplementing analysis by EEG-based regressors. This was even the case when the regressor that did reveal such differences when analyzing EEG alone. Next to left M1, when using these EEG-based regressors we also found positive correlations between source-reconstructed β-amplitudes in right M1, i.e. ipsilateral to the force producing hand. This was the case in both groups and does suggest the involvement of an ipsilateral controller in our unimanual perceptual motor task. Remarkably, the older group also showed a deactivation of a small part of the left M1. And, there were significant negative BOLD/ β-amplitude correlations in both groups in bilateral PM1 (as well as in the anterior intraparietal area and some visual areas). The latter activation patterns were only visible by combining fMRI and EEG. They do, indeed, confirm the idea of effective interhemispheric inhibition that we outlined in the *Introduction*. That is, contralateral M1 – in our case the left – projects to both the ipsilateral (right) PM1 and M1. As discussed in Chettouf et al. (2020), unimanual movements are considered a special case of bimanual ones. The direction and location of both inhibition and facilitation appears to depend on the motor task that is performed – an increase in task complexity seems to be accompanied with more (efficient) communication between hemispheres. When combining fMRI and EEG we do see bilateral cortical (de)activation in relevant motor areas during this uni-manual motor task in both groups. We expected that these interactions through the CC would be altered with age and lead to an imbalance of the effective interhemispheric inhibition. Yet, we do not find any significant age-related differences in this combined EEG-fMRI analysis despite the significant differences in EEG (Fig. 7). Perhaps this is because of (a) limited statistical power due to a small sample size per group (N = 13 versus 14), (b) both groups were able to learn the motor task to similar degree, and (c) the difference present in EEG has no principle correlate in fMRI – it is the degree of synchronization/desynchronization difference rather than of net synaptic activity.

A recent study with a similar design revealed the same behavioral results: a visuo-motor tracking skill was learned to a similar extent in both young and older adults (no interaction effect), while motor performance was lower in the older compared to the young (Berghuis et al., 2019). There older adults display stronger fMRI-BOLD activation. While this may seem to contradict our finding one should note that Berghuis et al. (2019) added the whole-brain gray matter volume a covariate to their model, the observed differences between younger and older adults were no longer significant. Apparently, differences in gray matter explained the age-related differences that were found, at least to some extent. In line with our findings, both groups showed a similar decrease of brain activation over time (from pre- to post-test), suggesting that the visual processing areas are more involved when performing the task for the first time compared to immediately after the training session. Yet, changes in brain-deactivations were age-dependent in specific areas that are part of the DMN (Raichle, 2015). This agrees with the idea that DMN modulation is ‘dysregulated; with increasing age (Park and Reuter-Lorenz, 2009) and suggests the employment of compensatory mechanisms by older adults to achieve similar learning rates as younger ones.

With respect to the postulated three alternative models of network interactions supporting sensorimotor coordination (Chettouf et al. (2020), Fig. 1) our present data support a significant age-related change of oscillatory activity in premotor areas in line with model B). Decreased capability of pre-motor β-rhythm desynchronization during the task leads to less efficient feedforward inhibition of ipsilateral M1 and to a decreased performance with increasing age. In subsequent studies, we will use multimodal data in the context of personalized brain network models that will further help us to disentangle the complex nonlinear network interactions underlying observed behavior (Ritter et al., 2013).

## 6. Conclusion

Demanding sensorimotor coordination performance that improves with learning is diminished in the older adults. We found that (1) Rolandic β-power in the pre-motor areas during learning is stronger (i.e. less task-related desynchronization) in younger than in older adults. And, (2) Rolandic β-power correlates negatively with fMRI in PM1 and positively in M1. Negative fMRI in PM1 during higher β-power and decreased β-power in PM1 during learning in older adults suggest a decreased PM1-mediated intra-hemispheric inhibition of M1 as a potential source for elevated interhemispheric crosstalk that may explain diminished motor performance at increased age.

## Acknowledgements

Part of the computation has been performed on the High Performance Computing for Research cluster of the Berlin Institute of Health. PR acknowledges support by EU H2020 Virtual Brain Cloud 826421, Human Brain Project SGA2 785907; Human Brain Project SGA3 945539, ERC Consolidator 683049; German Research Foundation SFB 1436 (project ID 425899996); SFB 1315 (project ID 327654276); SFB 936 (project ID 178316478; SFB-TRR 295 (project ID 424778381); SPP Computational Connectomics RI 2073/6-1, RI 2073/10-2, RI 2073/9-1; Berlin Institute of Health & Foundation Charité, Johanna Quandt Excellence Initiative.

## Supplementary material

### S.1 MR artifact correction EEG data

Combining the two modalities brings several technical challenges. The most prominent being the electromagnetic induction and the emergence of an electromotive force in a conductor enclosed by a change (Ritter and Villringer, 2006). Assuring measurements with minimal head movements and low impedance of the EEG channels is very important.

The EEG data were segmented into three segments: the experimental learning task, pre- and post-resting state. To detect the scanner gradient artifact, the data of every EEG-channel were notch filtered (band-with of 0.1 Hz) at the peak frequency of the scanner artifact (16.5 Hz) and its ten higher harmonics. Subsequently, the time series were split up into the length of single MR scans and underwent a principal component analysis. Components that contained the scanner artifact were detected and removed from the data whenever they displayed strong power at the aforementioned scanner frequency (or higher harmonics). A component was considered an artifact when at least ten peaks corresponded to the scanner artifact frequency of 16.5 Hz and exceeded two times the standard deviation of that principal component. To further smoothen the data and to increase the signal-to-noise ratio without distorting the signal, we finally applied a Savizky-Golay polynomial filter (1^st^ order, window size 21 samples ~ 4.2 ms).

We evaluated this gradient artifact correction of EEG data to assure the quality of the data using for further analysis (Ritter et al., 2007). To this end, we determined the ratio between the root-mean-square (RMS) of uncorrected gradient artifact EEG periods and corrected gradient-artifact periods. A high RMS-ratio indicates a strong reduction of the signal due to artifact elimination.

### S.2 EEG beamformer results for alpha-power

When comparing the motor task against rest, we found occipital and parietal activity in the α-band in both groups (*p* < .05), with a peak value in superior parietal cortex 7A (SPL 7A). This is part of Brodmann area 7, involved in vision and proprioception.

**Figure S1.**
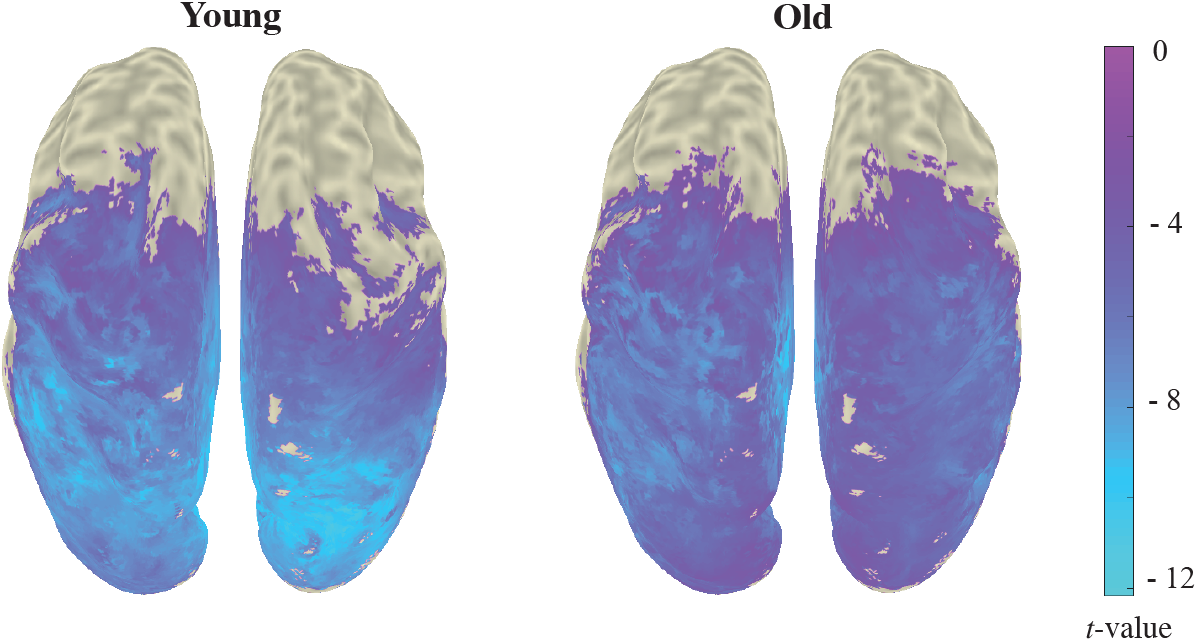
Mean power beamformer results in the α-band. Adaptive spatial filtering identified cortical sources where α-band power correlated when contrasting the motor task against baseline resting state. Grand averages shown correspond to the younger (N=13; left) and older (N=14, right) participants. Permutation tests revealed voxels comprising significant differences between task and rest on the group level. Colors represent *t*-values, masked with a threshold of *p* < .05.

